# Biochemical, biomarker, and behavioral characterization of the *Grn^R493X^* mouse model of frontotemporal dementia

**DOI:** 10.1101/2023.05.27.542495

**Authors:** Denise M. Smith, Geetika Aggarwal, Michael L. Niehoff, Spencer A. Jones, Subhashis Banerjee, Susan A. Farr, Andrew D. Nguyen

## Abstract

Heterozygous loss-of-function mutations in the progranulin gene (*GRN*) are a major cause of frontotemporal dementia due to progranulin haploinsufficiency; complete deficiency of progranulin causes neuronal ceroid lipofuscinosis. Several progranulin-deficient mouse models have been generated, including both knockout mice and knockin mice harboring a common patient mutation (R493X). However, the *Grn^R493X^* mouse model has not been characterized completely. Additionally, while homozygous *Grn^R493X^* and *Grn* knockout mice have been extensively studied, data from heterozygous mice is still limited. Here, we performed more in-depth characterization of heterozygous and homozygous *Grn^R493X^* knockin mice, which includes biochemical assessments, behavioral studies, and analysis of fluid biomarkers. In the brains of homozygous *Grn^R493X^* mice, we found increased phosphorylated TDP-43 along with increased expression of lysosomal genes, markers of microgliosis and astrogliosis, pro-inflammatory cytokines, and complement factors. Heterozygous *Grn^R493X^* mice did not have increased TDP-43 phosphorylation but did exhibit limited increases in lysosomal and inflammatory gene expression. Behavioral studies found social and emotional deficits in *Grn^R493X^* mice that mirror those observed in *Grn* knockout mouse models, as well as impairment in memory and executive function. Overall, the *Grn^R493X^* knockin mouse model closely phenocopies *Grn* knockout models. Lastly, in contrast to homozygous knockin mice, heterozygous *Grn^R493X^* mice do not have elevated levels of fluid biomarkers previously identified in humans, including neurofilament light chain (NfL) and glial fibrillary acidic protein (GFAP) in both plasma and CSF. These results may help to inform pre-clinical studies that use this *Grn* knockin mouse model and other *Grn* knockout models.

## INTRODUCTION

Progranulin is a widely expressed protein with pleiotropic effects, including growth factor-like properties, anti-inflammatory effects, promoting neurite outgrowth, and promoting autophagy [1-3]. However, the precise molecular function of progranulin is unclear. Within cells, progranulin localizes to lysosomes [4]; progranulin is also secreted from cells and found in blood and cerebrospinal fluid [5].

Heterozygous loss-of-function mutations in the progranulin gene (*GRN*) are a major cause of frontotemporal dementia (FTD) due to progranulin haploinsufficiency [6-8]. FTD involves atrophy of the frontal and temporal lobes, with onset usually occurring between 40 to 55 years of age. *GRN* mutations cause behavioral variant of FTD (bvFTD), which often presents with decline in social conduct, apathy, emotional blunting, and loss of insight [9]. Neuropathologically, FTD-*GRN* is characterized by cytoplasmic ubiquitin-positive inclusions of TAR DNA-binding protein 43 (TDP-43), neuroinflammation, and no tau pathology [6,7,10,11]. Several fluid biomarkers have been identified in the CSF and blood of individuals with FTD, including neurofilament light chain (NfL), glial fibrillary acidic protein (GFAP), TDP-43, and phosphorylated TDP-43 [12-18]. NfL is a particularly promising biomarker for FTD-*GRN*, because its levels correlate with disease severity [12,13]. Additionally, in asymptomatic *GRN* mutation carriers, high plasma NfL is predictive of disease conversion and progression during the subsequent two-year period [19].

In contrast to heterozygous *GRN* mutations, complete progranulin deficiency causes neuronal ceroid lipofuscinosis (NCL), CLN11, with disease onset usually in the teenage to midlife years, although this can vary widely [20-22]. NCL/CLN11 presents with vision loss, seizures, and ataxia. NCL is a form of lysosomal storage disease, with characteristic lipofuscin accumulation, pointing to an important function of progranulin for proper lysosome function and/or homeostasis. Moreover, FTD-*GRN* and NCL highlight the gene dosage effect of progranulin deficiency which manifests as distinct diseases.

Several progranulin-deficient mouse models have been generated by different labs, including four *Grn* knockout mouse lines [23-26] and two *Grn* knockin mouse lines that harbor a common FTD-associated nonsense mutation [27,28]. Overall, similar phenotypes have been described for all these mouse lines, although most reports have focused on homozygous mice. Homozygous *Grn* knockout and knockin mice exhibit signs of lysosomal impairment, including lipofuscin accumulation, NCL-like lysosomal storage material by electron microscopy, and upregulation of lysosomal genes in the brain which is likely to compensate for the lysosomal impairment [28-34]. Homozygous *Grn* knockout and knockin mice also show age-dependent neuroinflammation and cytoplasmic accumulation of phosphorylated TDP-43 [25,27,28,32,34-37]. Additionally, *Grn* knockout and *Grn^R493X^* knockin mice exhibit deficits in social and emotional behavior, including decreased sociability, age-dependent changes in social dominance, increased anxiety, and increased obsessive–compulsive behaviors [28,35-40]. Lastly, memory impairment has also been reported in *Grn* knockout mice, including in the Morris water maze and the novel object recognition test [24,36,40].

There are currently several important gaps in our knowledge about *Grn* mouse models. Most notably, characterization of heterozygous *Grn*^+/–^ and *Grn^+/R493X^* is not complete. For example, do these mice develop TDP-43 pathology, lysosomal dysfunction, neuroinflammation, or memory impairment, similar to the homozygous mice? Additionally, are the biomarkers identified in human *GRN* mutation carriers similarly increased in *Grn* mouse models? In the current study, we performed more in-depth characterization of heterozygous and homozygous *Grn^R493X^* knockin mice, which includes biochemical assessments, behavioral studies, and analysis of fluid biomarkers. These results may help to inform pre-clinical studies that use this and other *Grn* mouse models.

## MATERIAL AND METHODS

### Mice

*Grn^R493X^* knockin [28] and *Grn^–/–^* knockout mice [23] were on the C57BL/6J background (backcrossed more than 8 generations) and were genotyped by real-time PCR (Transnetyx). Mice were housed in a pathogen-free barrier facility with a 12-h light/12-h dark cycle and allowed food and water *ad libitum*. Experimental cohorts were generated by crossing heterozygous mice, and wild-type littermates were used for comparisons. All experimental procedures were approved by the Institutional Animal Care and Use Committee at Saint Louis University or the Harvard Medical Area Standing Committee on Animals.

### Experimental design

At the indicated ages, tissues were isolated, flash frozen in liquid nitrogen, and stored at -80 °C until analyzed. Blood was collected in EDTA-coated tubes (Sarstedt, Microvette CB 300 K2 EDTA, 16.444.100); tubes were centrifuged at 2000 × *g* for 5 min and the plasma was removed and stored at -80 °C. For the experiments of Figs. 4A and S2, blood was collected from the submandibular vein. In all other studies, trunk blood was collected. CSF was collected by capillary action from the fourth ventricle of anesthetized mice using a 30-gauge needle attached to Intramedic polyethylene tubing (Becton Dickinson, PE-10).

### Behavioral studies

A cohort of 11-month-old male wild-type, heterozygous, and homozygous *Grn^R493X^* mice underwent a test battery of behavioral tests to assess behavioral and cognitive function. Behavioral assessment was performed between 7am and 12pm by experimenters who were blinded to the genotypes of the mice. Mice were individually housed one week prior to the start of behavioral testing until the end of the study to ensure equal levels of arousal during behavioral testing. Each mouse received only one test per day. Following the behavioral studies, tissues were collected as described above at 12 months of age. For the tube test, an additional cohort of 8- to 11-month-old male mice tested by the NeuroBehavior Laboratory at the Harvard NeuroDiscovery Center was included to achieve appropriate statistical power.

### Open field test

The open field test was used to assess exploratory and locomotor activity. For this test, each mouse was allowed to freely roam an empty arena (53 cm × 63 cm) under dim white light (approximately 50 lux) for 5 min. The distance traveled and the time spent in the center of the open field were determined using the ANY-maze video tracking system (San Diego Instruments).

### Forced swim test

The forced swim test was used to assess depressive behavior. Each mouse was placed in an inescapable transparent tank (30 cm height × 20 cm diameter, with a water level of 15 cm) for 6 min. The time spent moving versus the time spent immobile was recorded using the ANY-maze tracking system. Analysis was performed by comparing the time spent immobile during the last 4 min between groups. The greater the amount of time spent immobile indicates increased depressive behavior. The water temperature was monitored and maintained at 25 °C during testing by exchanging cold water in the maze with warm tap water as needed. During testing, the mouse was continually monitored and rescued from drowning should it stop swimming or become compromised. After each trial, the mouse was placed in a cage with a warmed absorbent surface under a warming lamp until dry.

### Nesting behavior

Nest building was used to assess general well-being and motor systems [41]. At the start of the study, one 4-gram Bed-r’Nest white shredded paper disc (Andersons, BRN4WSR) was placed in the front of a fresh shoebox cage that contained the equivalent of 200 ml of corn cob bedding in the bottom, along with a fresh wire top containing food and water. Each mouse was placed in the cage in the morning at 9am and left for 24 h, and then the nest was photographed. Nests were scored on a 5-point scale: Score 1 = shredded paper was scattered throughout the cage or disc was untouched. Score 2 = some shredded material was used to construct the nest, but >50% of material was scattered and untouched. Score 3 = a noticeable nest was constructed, but several pieces of shredded paper were still scattered. Score 4 = almost all material used for nest, and few pieces remained scattered or near the nest. Score 5 = all material used to make an identifiable nest.

### Three-chamber sociability test

The three-chamber test [42] was used to assess the sociability of a mouse when given the choice between interacting with another mouse versus an inanimate object. For this test, each mouse was allowed to freely explore the empty three-chambered box (20.5 cm × 40.5 cm × 22 cm high) made of Plexiglas containing two empty wire cups in the outer chambers. After 10 min, a stranger mouse was placed under one of the wire cups and an inanimate object (a block) was placed under the other wire cup. During the next 10 min, the times spent interacting with the novel mouse and with the inanimate object were determined using the ANY-maze tracking system. Mice that did not interact with the novel mouse or the inanimate object were excluded from the data analysis. A sociability ratio was calculated for each mouse by dividing the time spent interacting with the novel mouse by the time spent interacting with the inanimate object.

### Tube test

The tube test was used to assess social dominance [38]. For this test, a wild-type mouse and a heterozygous *Grn^R493X^* mouse from separate home cages were paired; each mouse was tested against up to three mice of the other genotype. Mice were tested without habituation to the tube. Mice were placed, head first, into opposing sides of a clear vinyl tube (3.8 cm internal diameter × 30.5 cm length) and released simultaneously. The first mouse to place two paws outside of the tube was considered to be the less dominant mouse. If the mice crossed each other or no mouse left the tube after 2 min, the trial was aborted and later repeated at the end of the session. The winning percentage for each genotype was calculated based on n = 30 trials.

### Elevated plus maze

The elevated plus maze was used to assess anxiety and disinhibition-like behavior. This test used a plus-shaped apparatus with two open arms and two enclosed arms, which is elevated 50 cm above the floor. Each arm is 35.5 cm in length; two opposite arms are open while the other two opposite arms are enclosed, as previously described [43]. For this test, each mouse was placed on the central platform facing an enclosed arm and allowed to freely explore the maze for 5 min. The time spent in open arms was monitored using the ANY-maze tracking system.

### T-maze

The T-maze was used to assess long-term spatial memory [44] and involves the mouse learning to traverse the maze and turning in the correct direction to find the goal box. The maze consists of a black plastic alley with a start box at one end and two goal boxes at the other. The start box is separated from the alley by a plastic guillotine door that prevents movement down the alley until raised at the onset of training. An electrifiable floor of stainless-steel rods run throughout the maze to deliver a mild scrambled foot-shock 5 sec after the guillotine door is raised and a cue buzzer (door-bell type sounded at 55 dB). For this test, each mouse was trained to one avoidance (of a mild 0.35 mA foot shock). One week later, memory retention was tested, with a criterion for retention defined as 5 avoidances in 6 consecutive trials.

### Y-maze

The Y-maze was used to assess working memory [45]. This test relies on the innate curiosity of rodents to explore previously unvisited areas, and it measures the number of visits to consecutive arms without revisiting the arm just visited. For this test, each mouse was placed in a Y-shaped maze for 5 min and the sequence of arm visits was monitored using the ANY-maze tracking system. The percentage of arm visits was determined as % alternations = (number of alternations / total number of arm entries) x 100.

### Novel object recognition

The novel object recognition test was used to assess short-term memory [46]. This test relies on the innate curiosity mice to explore novel things in their environment, and it measures the ability of the mice to recognize the previously explored object. For this test, each mouse was allowed to explore a maze with two like objects placed on one wall for 5 min. One hour after the initial phase, the mouse was placed in the maze with one like object in its original place and one new object on the opposite wall for 5 min. The time spent exploring the old object and the new object was recorded, and the discrimination index (DI) was calculated as DI = (T_new_ – T_old_) / (T_old_ + T_new_).

### Puzzle box

The puzzle box was used to assess complex problem solving and memory [47,48]. This test relies on the innate desire of mice to avoid open spaces, and it measures latency to find the dark side of the maze by traversing a tunnel with increasingly difficult obstacles placed at the opening of the tunnel. The puzzle box is an arena consisting of a Plexiglas white box divided by a removable barrier into two compartments: a brightly lit start zone (58 cm × 28 cm) and a smaller, dark goal zone (15 cm × 28 cm). For this test, each mouse was introduced into the start zone and trained to move into the goal zone through a narrow underpass (∼ 4 cm wide) located under the barrier. Each mouse underwent a total of 11 trials over 4 consecutive days, during which they were challenged with obstructions of increasing difficulty placed at the tunnel entrance: completely opened (Trials 1-2), partially blocked by a guillotine (Trials 3-4), blocked by nesting paper (Trials 5-7), or blocked by a T-shaped cardboard plug (Trials 8-11). Mice that took > 4 min to complete the first 3 trials or that refused to perform the task for more than one trial were excluded from the data analysis. This experimental paradigm allows for assessment of problem solving ability, learning/short-term memory, and long-term memory [47].

### qPCR

Total RNA was isolated from flash frozen tissues using the RNeasy Mini kit (Qiagen, 74106) with on-column DNase digestion (Qiagen, 79256). RNA was reverse-transcribed to obtain cDNA using the iScript cDNA synthesis kit (Bio-Rad, 1708891), and qPCR was performed using PowerUp SYBR Green Master Mix (ThermoFisher, A25777) with a Bio-Rad CFX384 Real-Time System. Primers sequences are provided in Table S1. Results for qPCR were normalized to the average of two housekeeping genes (*36B4* and *Cyclo*) and analyzed by the comparative C_T_ method.

### Western blot

Cortical tissues were homogenized and fractionated as previously described [30] in buffers containing protease inhibitors (Roche, cOmplete Mini EDTA-free Protease Inhibitor Cocktail) and phosphatase inhibitors (Roche, PhosSTOP). Protein concentrations were determined for the high salt (HS) and RIPA fractions using the Bio-Rad DC Protein Assay Kit II. For western blot analysis, sample buffer was added to the lysates, and the samples were heated at 95 °C for 10 min. Equal amounts of protein lysates were separated on SDS–PAGE gels. Proteins were transferred to nitrocellulose membranes using the Bio-Rad Turbo-Blot transfer system. After blocking and antibody incubations, membranes were incubated with SuperSignal West Pico or Femto enhanced chemiluminescent HRP substrate (ThermoFisher) and visualized using a Chemi-Doc system (Bio-Rad). Primary antibodies used for immunoblot analysis include: an anti-mouse progranulin polyclonal antibody [49] used at a 1:3000 dilution, an anti-cathepsin D monoclonal antibody (abcam, ab75852, 1:500), an anti-cathepsin L polyclonal antibody (Cell Signaling Technology, 71298S, 1:500), an anti-LAMP1 monoclonal antibody (Developmental Studies Hybridoma Bank, 1D4B, 1:500), an anti-β-actin monoclonal antibody (Cell Signaling Technology, 4970S, 1:3000), an anti-Iba1 polyclonal antibody (abcam, ab5076, 1:500), an anti-GFAP polyclonal antibody (abcam, ab134436, 1:1000), an anti-TDP-43 polyclonal antibody (Proteintech, 10782-2-AP, 1:2000), and an anti-phosphorylated TDP-43 (Ser409/410) monoclonal antibody (BioLegend, 829901, 1:250). HRP-conjugated secondary antibodies were from Jackson Immuno Research Labs: goat anti-rabbit IgG (111035144), donkey anti-rat IgG (712035150), donkey anti-goat IgG (705035147), and donkey anti-chicken IgG (703035155). Data were quantified by densitometry using ImageJ and normalized to β-actin levels in the respective fractions.

### ELISA

C1qa and C3 levels were measured using ELISA kits from abcam (ab291069 and ab26388) according to the manufacturer’s instructions. In Fig. 2C, 10 µg of cortical lysates were used. In Fig. 2D, CSF samples were diluted 500-fold.

### Biomarkers

NfL and GFAP levels were measured using the Quanterix Simoa HD-1 platform with the Simoa NF-L Advantage Kit or the Neurology 2-Plex B Advantage Kit, respectively. Plasma and CSF samples were diluted 40-fold and 200-fold, respectively. All samples were assayed in duplicate wells. For a subset of plasma samples, NfL levels were also measured using the Sigma Single Molecule Counting (SMC) platform with the SMC NF-L High Sensitivity Immunoassay Kit, with plasma samples diluted 16-fold.

### Statistical analyses

Data are presented as mean ± SEM and were analyzed with GraphPad Prism software. Most comparisons were analyzed by one-way ANOVA with Dunnett post hoc test. In the experiment of Figs. 4A and S3A, data were analyzed by repeated measures two-way ANOVA with Tukey post hoc test. In the experiment of Fig. 5B, the distribution of wins per mouse was analyzed by two-way binomial test, with the expected distribution set at 50% wins for each genotype, similar to what has been previously described [38]. P values < 0.05 were considered significant.

## RESULTS

### Changes in lysosomal gene expression, inflammation, and TDP-43 levels in the brains of Grn^R493X^ mice

Previous studies of *Grn* knockout mouse models have documented increased expression of lysosomal proteins in the brain [29-32], likely to compensate for impaired lysosomal function. To investigate if *Grn^R493X^* mice exhibit similar changes, we analyzed mRNA and protein levels in brain tissues from *Grn^R493X^* mice at 6 months and 12 months of age. As expected, heterozygous and homozygous knockin mice had decreased *Grn* mRNA and progranulin protein levels in the cortex (Fig. 1). Heterozygous and homozygous *Grn^R493X^* mice both showed modest increases in expression of lysosomal proteins (including TFEB, CtsD, PSAP, and LAMP1) in the cortex at 6 months and 12 months of age (Fig. 1A). Similar changes were also observed in the thalamus of *Grn^R493X^* mice (Fig. S1). At the protein level, increased LAMP1 levels were observed in the cortex of 12-month-old heterozygous and homozygous *Grn^R493X^* mice (Fig. 1B and Fig. S2). Increased CTSD levels were limited to homozygous mice. For CTSD, these increases were observed in both the proprotein and the cleaved mature form. These results indicate that *Grn^R493X^* mice exhibit a broad increase in lysosomal gene expression in the brain, similar to *Grn* knockout mice.

**Figure 1.**
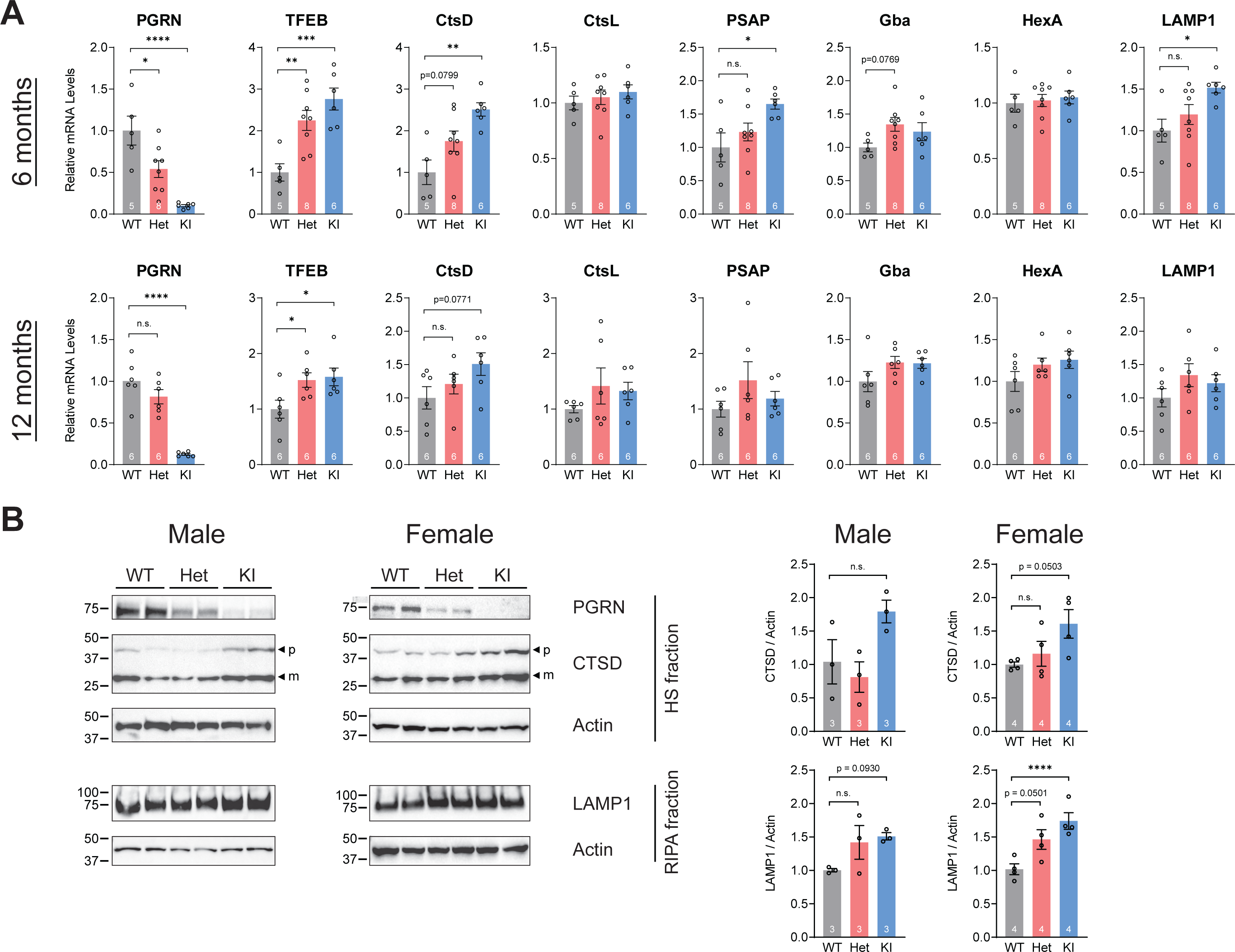
*Grn^R493X^* mice have increased expression of multiple lysosomal genes in the cortex. A) Quantification of mRNA levels in the cortex by qPCR (male and female mice). B) Western blot analysis of selected lysosomal proteins in the cortex of 12-month-old mice (males and females, as indicated). Arrowheads indicate the proprotein (p) and mature cleaved (m) forms of CTSD. Bars represent mean ± SEM; the group sizes are indicated in each bar. * p < 0.05, ** p < 0.01, *** p < 0.001, and **** p < 0.0001 compared to wild-type group, as determined by one-way ANOVA with Dunnett post hoc test. WT, wild-type; Het, *Grn^+/R493X^* heterozygous mice; KI, *Grn^R493X/R493X^* knockin mice; n.s., not significant.

Age-dependent neuroinflammation is a major phenotype that has been observed in multiple *Grn* knockout mouse models [23,25,29,32,35], although examination in heterozygous mice has been more limited [30,34,35]. For the *Grn^R493X^* mouse model, we previously reported age-dependent microgliosis in the thalamus of 12-month-old homozygous mice, as determined by quantification of Iba1^+^ cells after immunohistochemistry [28]. Additionally, Frew and Nygaard reported increased microgliosis and astrogliosis in 18-month-old homozygous mice by immunostaining [37]. Here, we report increased expression of the microglia marker Iba1 and astrocyte marker GFAP in the cortex that is apparent in homozygous mice by 12 months of age (Fig. 2A). In contrast, heterozygous *Grn^R493X^* mice did not show a significant increase. Western blot analysis of cortical tissue from 12-month-old mice confirmed the increase in GFAP at the protein level in homozygous mice (Fig. 2B and Fig. S2). Iba1 levels were significantly increased in homozygous female mice, but not in homozygous male mice. Overall, these observations in *Grn^R493X^* mice are consistent with a previous report in 12-month-old *Grn* knockout mice [35].

**Figure 2.**
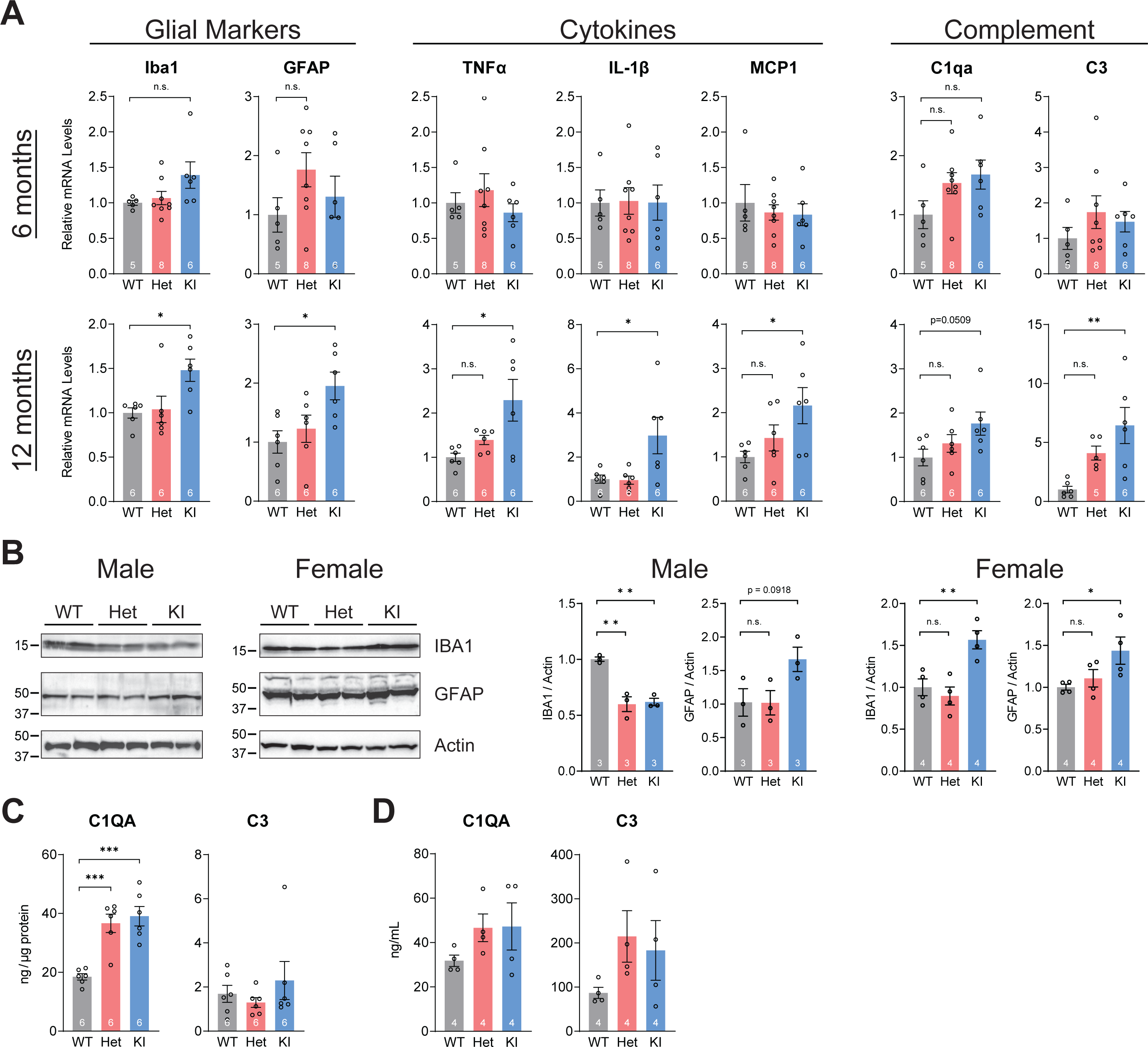
*Grn^R493X^* mice exhibit age-dependent neuroinflammation in the cortex and CSF. A) Quantification of mRNA levels by qPCR (male and female mice). B) Western blot analysis of selected inflammation-related proteins in 12-month-old mice (males and females, as indicated). C-D) C1qa and C3 protein levels in cortical tissue (C) and CSF (D) of 12-month-old female mice, as determined by ELISA. Bars represent mean ± SEM; the group sizes are indicated in each bar. * p < 0.05, ** p < 0.01, and *** p < 0.001 compared to wild-type group, as determined by one-way ANOVA with Dunnett post hoc test. WT, wild-type; Het, *Grn^+/R493X^* heterozygous mice; KI, *Grn^R493X/R493X^* knockin mice; n.s., not significant.

We found modest increases in expression of cytokines (including TNFα, IL-1β, and MCP1) in the cortex of homozygous *Grn^R493X^* mice at 12 months of age (Fig. 2A). Heterozygous mice had intermediate cytokine levels, which were not statistically different than the wild-type mice. Notably, we observed a marked increase in the expression of complement factor C3 in 12-month-old homozygous mice; trends toward increased levels of C1qa and C3 were also seen in heterozygous *Grn^R493X^* mice, although the magnitude of the increases was smaller than in homozygous mice. At the protein level, C1qa levels were significantly increased in the cortex of both heterozygous and homozygous *Grn^R493X^* mice (Fig. 2C). Additionally, in CSF, C1qa and C3 levels trended higher in heterozygous and homozygous *Grn^R493X^* mice (Fig. 2D); however, these differences were not significant, possibly due to the small sample sizes. Together, these results indicate that *Grn^R493X^* mice develop age-dependent neuroinflammation that is similar to previous reports in *Grn* knockout mouse models.

TDP-43 pathology is a hallmark feature of FTD-*GRN* [10,50]. We and others have reported increased cytoplasmic and phosphorylated TDP-43 in the brains of homozygous *Grn* knockout mice [30,51,52] and homozygous *Grn^R493X^* knockin mice [27,28]. A study by Götzl et al. showed that heterozygous *Grn*^+/–^ mice do not have increase in phosphorylated TDP-43 levels in the brain [30]. Here, we measured levels of total and phoshphorylated TDP-43 (Ser409/410) in the RIPA-soluble and insoluble fractions of cortical tisuses of 12-month-old *Grn^R493X^* mice. Compared to wild-type mice, homozygous female *Grn^R493X^* mice had decreased levels of soluble TDP-43 in the RIPA fractions (Fig. 3A and Fig. S2), which is similar to a previous report in 18-month-old homozygous *Grn^R493X^* mice [37]. Levels of insoluble TDP-43 were not significantly different in heterozygous and homozygous *Grn^R493X^* mice, compared to wild-type mice. In the insoluble fractions, we found increased levels of phosphorylated TDP-43 in homozygous female *Grn^R493X^* mice.

**Figure. 3.**
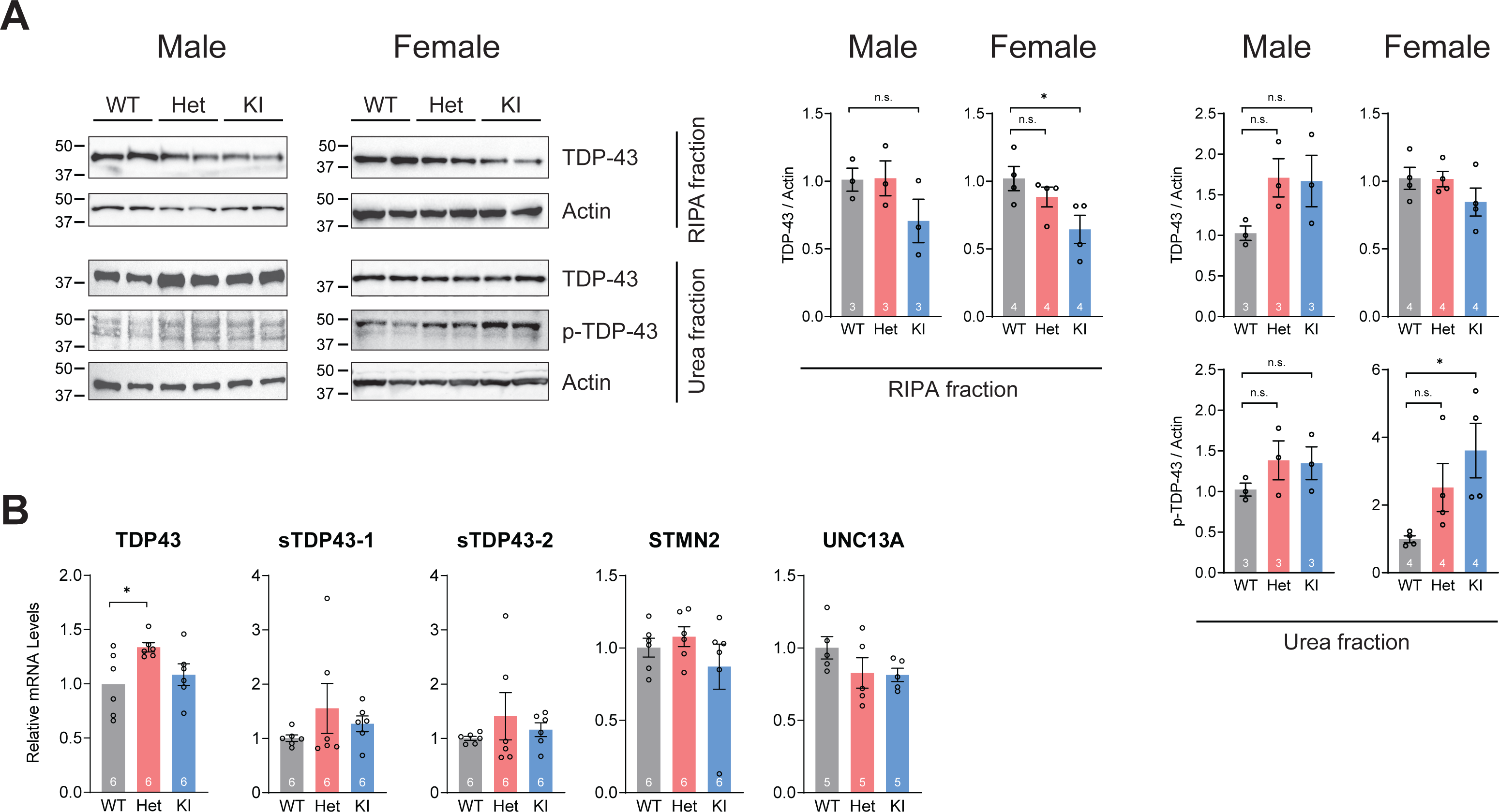
Homozygous *Grn^R493X^* mice have increased phosphorylated TDP-43 at 12 months of age. A) Western blot analysis of total and phosphorylated TDP-43 (Ser409/410) in the high salt (HS), RIPA-soluble, and RIPA-insoluble fractions of cortical tissue (males and females, as indicated). B) Quantification of mRNA levels by qPCR (male and female mice). Bars represent mean ± SEM; the group sizes are indicated in each bar. * p < 0.05 compared to wild-type group, as determined by one-way ANOVA with Dunnett post hoc test. WT, wild-type; Het, *Grn^+/R493X^* heterozygous mice; KI, *Grn^R493X/R493X^* knockin mice.

At the mRNA level, expression of TDP-43 was not increased in homozygous mice (Fig. 3B), although a modest increase was observed in the cortex of 12-month-old heterozygous mice. No differences were found in the levels of shortened TDP-43 splice isoforms, sTDP-43-1 and sTPD-43-2 [53]. TDP-43 functions to repress cryptic exon inclusion during RNA splicing, and TDP-43 depletion has been shown to perturb RNA metabolism [54]. We therefore measured levels of several mRNAs known to be regulated by TDP-43, including Stathmin-2 (STMN2) and Unc-13 homolog A (UNC13A); total levels of these mRNAs are decreased by TDP-43 depletion [55-58]. As shown in Fig. 3B, we found no significant differences in the levels of these mRNAs between genotypes. Together, these data indicate that heterozygous and homozygous *Grn^R493X^* mice have a modest increase in phosphorylated TDP-43 and a male-specific increase in insoluble TDP-43 in the cortex, but they do not have detectable defects in TDP-43-dependent RNA splicing.

### Biomarker levels in the Grn^R493X^ mouse model

Previous human studies have identified several biomarkers that are increased in the blood or CSF of individuals with FTD, including those with heterozygous *GRN* mutations. These biomarkers include NfL [12-14], GFAP [14,18], total TDP-43 [16], and phosphorylated TDP-43 [15,17]. In particular, NfL is a neuronal cytoskeletal protein that is released into CSF and blood upon neuronal damage or death. CSF and blood NfL levels have been proposed as an early indicators of neurodegeneration [59], and their levels reflect FTD disease severity [12,13]. It is currently not known if levels of NfL and other biomarkers are similarly increased in *Grn* knockout and *Grn^R493X^* mouse models. To address this, we measured plasma NfL levels in a cohort of *Grn^R493X^* mice between 2 months and 18 months of age using the Single Molecule Array (Simoa) platform by Quanterix. As shown in Fig. 4A, we observed an age-dependent increase in plasma NfL levels with all genotypes. We detected elevated levels in homozygous *Grn^R493X^* mice beginning at 10-12 months of age. In contrast, plasma NfL levels did not differ between wild-type and heterozygous *Grn^R493X^* mice. These trends were also observed when male and female mice were analyzed separately (Fig. S3). These trends were also observed with NfL measured using the Single Molecule Counting (SMC) platform by Sigma (Fig. S4); there was a strong correlation between the measurements from two assays in the 14-month-old plasma samples (r^2^ = 0.8541, n=23, p < 0.0001 by two-tailed t test). We measured plasma NfL levels in additional cohorts of 12-month-old and 18-month-old mice. While homozygous *Grn^R493X^* mice had increased NfL levels at both ages (Fig.4B), we again found no differences between wild-type and heterozygous *Grn^R493X^* mice at 12 months of age. In 18-month-old mice, we observed a trend toward increased plasma NfL levels in heterozygous *Grn^R493X^* mice, although this did not reach statistical significance (p = 0.1105). For comparison, we measured plasma NfL levels in the *Grn* knockout mouse model [23] at 12 and 18 months of age; similarly, we found increased NfL levels in homozygous *Grn* knockout mice but not in heterozygous mice (Fig. S5). We also measured NfL levels in a limited number of CSF samples; these results show elevated NfL levels in homozygous *Grn^R493X^* mice but not in heterozygous mice at 12 months of age (Fig. 4C).

**Figure 4.**
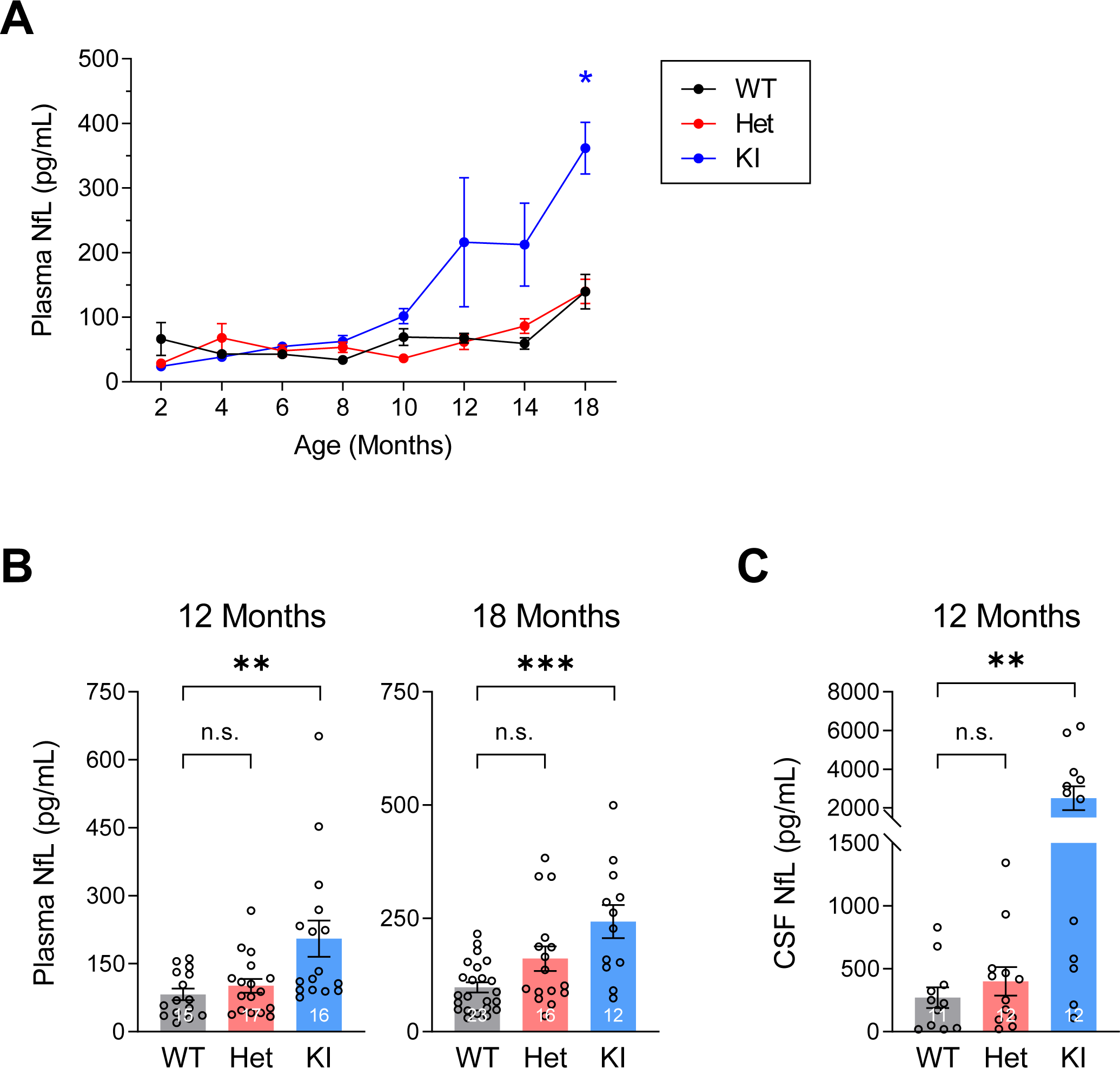
Homozygous *Grn^R493X^* mice, but not heterozygous mice, have increased NfL levels in blood and CSF. A) Plasma NfL levels in an aging cohort of mice (n = 6 per group, male and female mice). B) Plasma NfL levels in additional cohorts of mice (male and female mice). C) CSF NfL levels (male and female mice). Bars represent mean ± SEM; the group sizes are indicated in each bar. * p < 0.05, ** p < 0.01, and *** p < 0.001 compared to wild-type group, as determined by repeated measures two-way ANOVA with Tukey post hoc test (A) or one-way ANOVA with Dunnett post hoc test (B-C). WT, wild-type; Het, *Grn^+/R493X^* heterozygous mice; KI, *Grn^R493X/R493X^* knockin mice; n.s., not significant.

We also measured levels of another biomarker, GFAP, in the plasma and CSF of *Grn^R493X^* mice. In plasma, GFAP levels were not increased in heterozygous or homozygous *Grn^R493X^* mice at 12 months of age (Fig. S6A). In CSF, we observed a trend toward increased GFAP levels in homozygous *Grn^R493X^* mice but no increase in heterozygous mice at 12 months of age (Fig. S6B). Collectively, these results suggest that the FTD fluid biomarkers identified in humans are not similarly elevated in the blood and CSF of heterozygous *Grn*^+/–^ and *Grn^+/R493X^* mouse models. Additionally, the NfL results in particular suggest that, compared to their wild-type littermate controls, heterozygous *Grn^R493X^* mice do not exhibit increased neuronal death that is detectable by plasma or CSF NfL through 18 months of age.

As mentioned above, we also measured C1qa and C3 levels in CSF by ELISA in a small number of samples. Although the differences were not statistically significant, we observed trends towards increased levels of both C1qa and C3 in CSF of heterozygous and homozygous *Grn^R493X^* mice (Fig. 2D).

### Behavioral deficits in the Grn^R493X^ mouse model

Limited behavioral studies have been performed on the *Grn^R493X^* mouse model, and no data is currently available for heterozygous mice. Specifically, we have reported excessive grooming behavior, which is a type of obsessive compulsive-like behavior, in the homozygous *Grn^R493X^* mice [28], and Frew and Nygaard have reported increased anxiety in homozygous male mice [37]. Here, we performed additional behavioral studies in 11-month-old male mice to further characterize *Grn^R493X^* mice. Similar to previous reports in *Grn* knockout mouse models [35,36], we observed decreased sociability in homozygous *Grn^R493X^* mice in the three-chamber test (Fig. 5A). A similar trend was observed in heterozygous mice, although the difference was not statistically significant (p = 0.0869). In the tube test of social dominance, heterozygous *Grn^R493X^* mice had a lower winning percentage against wild-type mice (Fig. 5B). Although this difference was not statistically significant (p = 0.2005), it is suggestive of decreased social dominance, which would be in agreement with previous reports with similarly aged *Grn^+/–^* mice [38,60]. In the elevated plus maze, homozygous *Grn^R493X^* mice spent less time spent in the open arms, suggestive of increased anxiety (Fig. 5C). This finding is consistent with a previous study in homozygous *Grn* knockout mice using the elevated plus maze [61], as well as reports of increased anxiety in homozygous *Grn^R493X^* male mice [37] and in homozygous *Grn* knockout mice [26,24] based on time spent in the center of the open field. We observed no differences between genotypes in other behavioral tests, including open field activity, time spent in the center of the open field, forced swim test, and nest building (Fig. S7). Together, these studies show that *Grn^R493X^* mice exhibit social and emotional deficits similar to those previously reported in *Grn* knockout mouse models.

**Figure 5.**
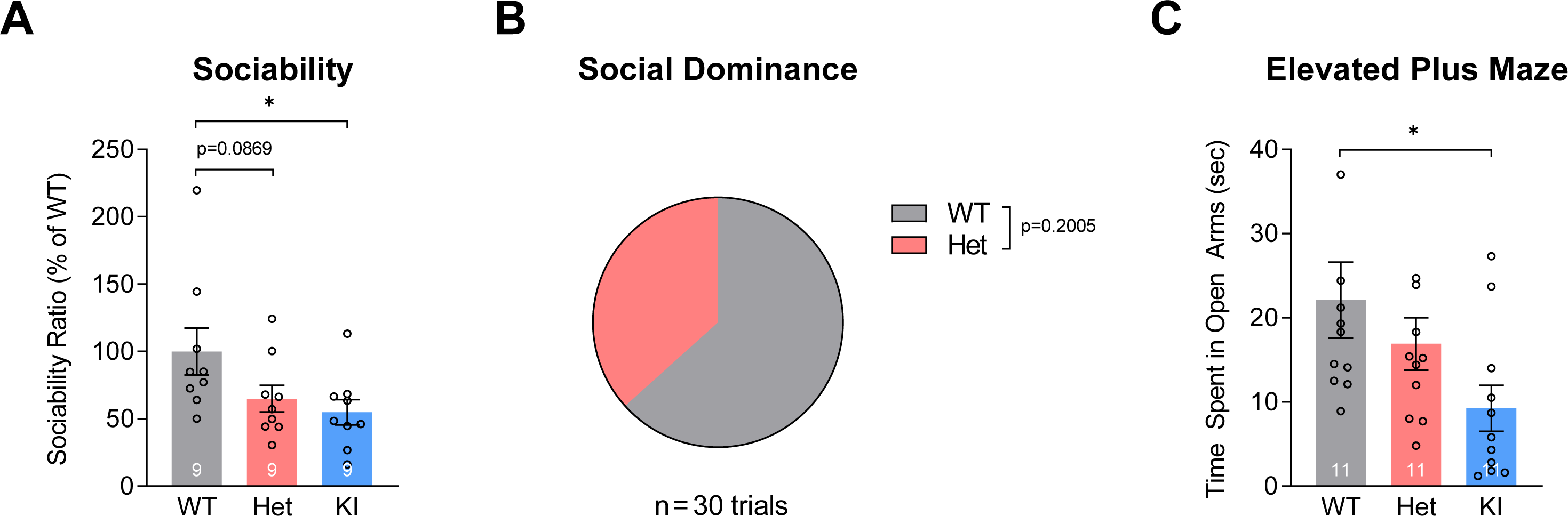
*Grn^R493X^* mice exhibit social and emotional deficits. A) Three-chamber test of sociability ( male mice, 11 months old). B) Tube test of social dominance (n = 15 per group, male mice, 8-11 months old). Percent of trials won is shown. C) Elevated plus maze (male mice, 11 months old). Bars represent mean ± SEM; the group sizes are indicated in each bar. * p < 0.05 compared to wild-type group, as determined by one-way ANOVA with Dunnett post hoc test (in A and C). Results were analyzed by two-way binomial test (in B). WT, wild-type; Het, *Grn^+/R493X^* heterozygous mice; KI, *Grn^R493X/R493X^* knockin mice.

Lastly, we assesed cognitive function in the *Grn^R493X^* mice. Using the T-maze to assess long-term spatial memory, we found impairment in heterozygous and homozygous *Grn^R493X^* mice (Fig. 6A). This impairment was refected in the increased number of trials needed to reach the criterion of choosing the correct side of the T-maze 5 out of 6 consecutive times. Using the novel object recognition test (with 1 h retention interval), heterozygous and homozygous *Grn^R493X^* mice exhibited impairment in short-term memory, as reflected in the lower discrimation index which is a calculation of the time spent interacting with the novel object (Fig. 6B). *Grn^R493X^* mice showed no differences in working memory, as assessed by the Y-maze (Fig. 6C). In the puzzle box test, we found impaired executive function in homozygous *Grn^R493X^* mice, but not in the heterozygous mice (Fig. 6D). Together, these results indicate that *Grn^R493X^* mice exhibit cognitive impairment in several specific domains. These results are consistent with previous reports of memory impairment in *Grn* knockout mice as assessed by the Morris water maze [36], novel object recognition test [24], and fear conditioning test [35].

**Figure 6.**
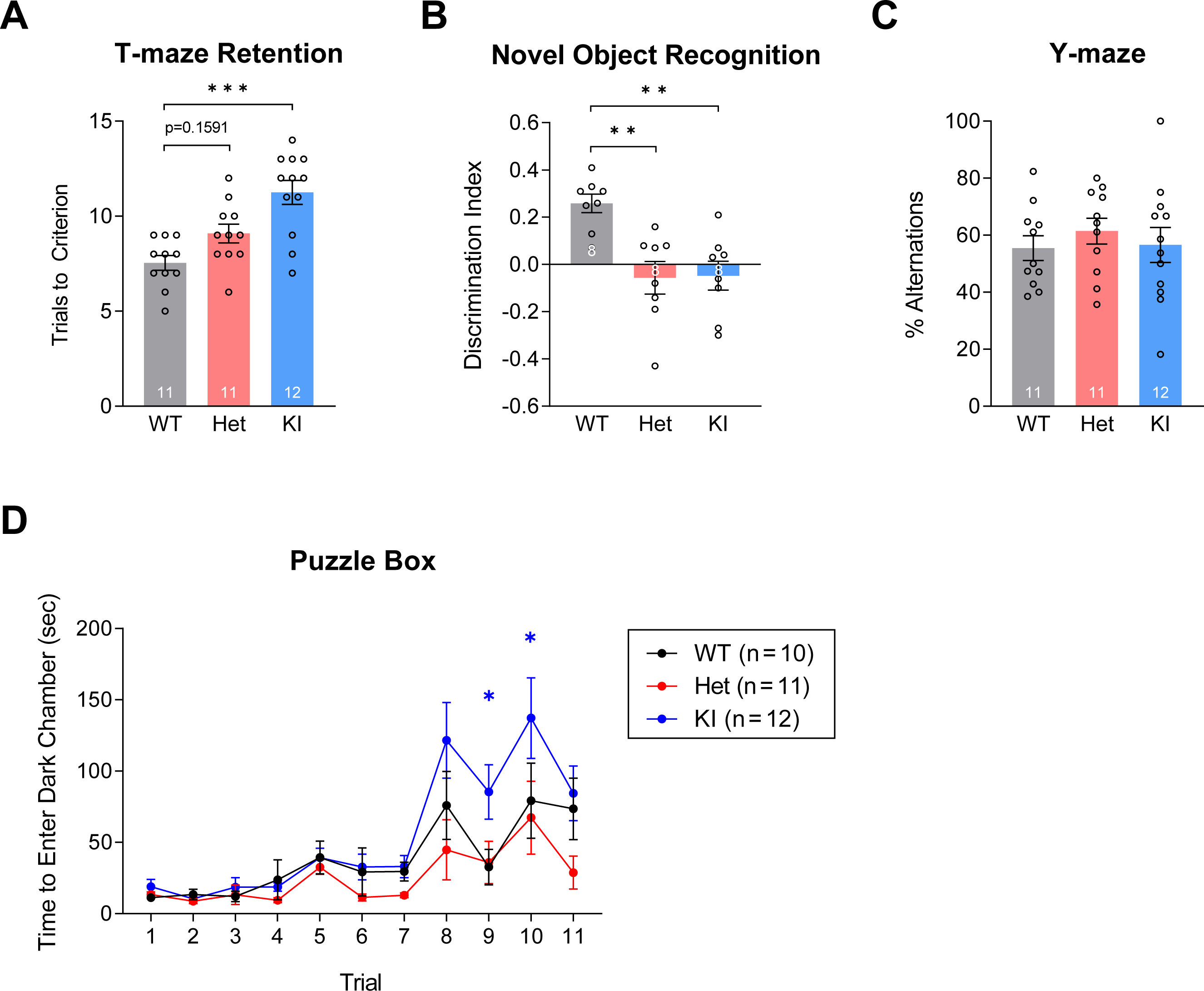
*Grn^R493X^* mice exhibit memory impairment at 11 months of age. A) T-maze retention (male mice). B) Novel object recognition with 1 h retention interval ( male mice). C) Y-maze (male mice). D) Puzzle box (male mice). Bars represent mean ± SEM; the group sizes are indicated in each bar. * p < 0.05, ** p < 0.01, and *** p < 0.001 compared to wild-type group, as determined by one-way ANOVA with Dunnett post hoc test. WT, wild-type; Het, *Grn^+/R493X^* heterozygous mice; KI, *Grn^R493X/R493X^* knockin mice.

## DISCUSSION

In the current study, we have further characterized the *Grn^R493X^* knockin mouse model of FTD, including analyses of biochemical changes, neuroinflammation, fluid biomarkers, and behavior. The R493X mutation introduces a premature termination codon, which leads to degradation of the mutant *Grn* mRNA through the nonsense-mediated RNA decay pathway, and therefore *Grn^R493X^* mice have markedly reduced progranulin protein levels [28]. Overall, our results show that *Grn^R493X^* knockin mice phenocopy their *Grn* knockout counterparts, with a mild behavioral phenotype and modest biochemical changes.

In the brain, heterozygous and homozygous *Grn^R493X^* mice had increased expression of lysosomal genes by 6 months of age (Figs. 1 and S1), likely to compensate for impaired lysosomal function and/or homeostasis. Similar increases in lysosomal gene expression have been reported in *Grn* knockout mice [29-32]. Additionally, lipofuscin has been observed in the brains of both homozygous *Grn*^–^*^/^*^–^ and *Grn^R493X^* mice [24,28,29,34,37,60,62], further indicating that progranulin deficiency leads to lysosomal dysfunction in these mouse models. It is unclear why the upregulation of some lysosomal genes (i.e., LAMP1, Gba, HexA, PSAP) was diminished at 12 months of age. Whereas the changes in lysosomal gene expression were evident at 6 months of age, increased expression of inflammation-related genes was generally not observed in *Grn^R493X^* mice until 12 months of age (Fig. 2). These genes include markers of microgliosis and astrogliosis, pro-inflammatory cytokines, and complement factors. Aside from C1qa and C3, changes in inflammation-related genes were largely absent in heterozygous *Grn^R493X^* mice. Overall, this pattern of neuroinflammation is consistent with previous reports in *Grn* knockout mouse models [25,29,34-36]. Together, these results suggest that neuroinflammation may be secondary to lysosomal dysfunction in the *Grn^R493X^* mouse model. TDP-43 pathology is a hallmark feature of FTD-*GRN* [10,50]. Previous studies have found increased cytoplasmic and phosphorylated TDP-43 in the brains of homozygous *Grn* knockout mice [30,51,52] and homozygous *Grn^R493X^* knockin mice [27,28], but effects have not been observed in heterozygous *Grn*^+/-^ mice [30]. In the current study, we did not find significant increases in phosphorylated TDP-43 (Ser409/410) levels in the cortex of 12-month-old heterozygous *Grn^R493X^* mice (Fig. 3A). Additionally, we did not observe effects on the levels of insoluble TDP-43 in the cortex of heterozygous or homozygous *Grn^R493X^* mice. Overall, these results are consistent with findings in other *Grn* knockout mouse models [30,51,52].

Our behavioral studies found social and emotional deficits in *Grn^R493X^* mice which mirror those found in *Grn* mouse models. Decreased sociability has been reported in both heterozygous and homozygous *Grn* knockout mice [35,36], and we similarly observed this in *Grn^R493X^* mice. Decreased social dominance has also been reported in 9- to 16-month-old heterozygous *Grn*^+/-^ mice [38,60], and our results suggest that heterozygous *Grn^+/R493X^* mice also exhibit this phenotype at 11 months of age; additional studies with larger cohorts are needed. A male-specific increase in anxiety has been reported in homozygous *Grn* knockout mice and homozygous *Grn^R493X^* mice [26,37,61]. In the current study, we found evidence of increased anxiety in homozygous *Grn^R493X^* male mice in the elevated plus maze (Fig. 5C), but we did not detect differences based on time spent in the center of the open field (Fig. S7B). It is possible that this discrepancy may reflect differences in these tests, such as the inclusion of both passive and active avoidance and approach/avoidance conflict in the elevated plus maze [63]. Indeed, it has been suggested that these differences may enable the elevated plus maze to detect anxiety-related changes that are not detected in other tests [64]. Additionally, similar to *Grn* knockout mice [39], we previously reported that homozygous *Grn^R493X^* mice have increased grooming behavior, which is a type of obssesive compulsive-like behavior in mice [28]. Together, our studies show that *Grn^R493X^* mice phenocopy *Grn* knockout mice with respect to behavior and that they mirror symptoms of FTD-*GRN* including diminished social interest, apathy, disinhibition, obsessive-compulsive behavior, and anxiety [9,65].

Memory is typically spared in bvFTD until later the stages of disease, although there are reports of memory impairment [9,66]. Three studies have reported memory impairment in homozygous *Grn* knockout mice using the Morris water maze, novel object recognition test, and fear conditioning test [24,35,36]. In the current study, we found that both homozygous *Grn^R493X^* mice as well as heterozygous mice show impariments in certain types of memory. Specifically, short-term memory and long-term spatial memory are impaired, but working memory was unaffected. In bvFTD, there is often impairment of executive functions which correlates with atrophy in the dorsolateral and medial prefrontal cortex [67,68]. Therefore, we assessed executive function in *Grn^R493X^* mice using the puzzle box test, which relies heavily on the prefrontal cortex as well as the hippocampus [47]. Overall, we found that homozygous, but not heterozygous, *Grn^R493X^* mice exhibit impaired executive function. These results show that 11-month-old heterozygous and homozygous *Grn^R493X^* mice have impairments in specific types of memory, and homozygous mice also have impaired executive function. This supports a previous study finding impaired associative learning and memory in the heterozygous *Grn*^+/-^ mice [35].

We sought to test if fluid biomarkers identified in individuals with FTD-*GRN* are similarly elevated in *Grn* knockout and *Grn^R493X^* mouse models. We largely focused on NfL, which is a neuronal cytoskeletal protein that has emerged as a promising biomarker that could be used to monitor the progression of neurodegenerative diseases and the effects of disease-modifying treatments [59]. While we observed increased NfL levels in the plasma and CSF of homozygous *Grn^R493X^* mice by 10-12 months of age, we did not find increased levels in heterozygous *Grn^R493X^* mice through 18 months of age (Figs. 4, S2, and S3). We also measured plasma NfL levels in the *Grn* knockout mouse model and similarly found elevated levels in homozygous but not heterozygous mice (Fig. S5). Examining levels of another FTD-*GRN* biomarker, we similarly found increased GFAP levels in the CSF of homozygous *Grn^R493X^* mice at 12 months of age, but no increase in heterozygous *Grn^R493X^* mice (Fig. S6C). Overall, these biomarker results suggest the absence of overt neuronal death in heterozygous *Grn*^+/-^ and *Grn^+/R493X^* mice. This contrasts the the severe atrophy of the frontal and temporal lobes observed in humans with heterozygous *GRN* mutations, which is accompanied by increases in NfL and GFAP in the CSF and plasma. The reasons for these differences between humans and mouse models are unclear. Nonetheless, they suggest that blood NfL and GFAP levels are not useful as biomarkers for pre-clinical studies using the heterozygous *Grn*^+/-^ and *Grn^+/R493X^* mouse models. Thus, it will be worthwhile to validate alternative blood biomarkers, such as compelement C1qa and C3 [32,69] or possibly specific lipid species such as gangliosides [70,71] in these mouse models. Indeed, our limited analysis suggests C1qa and C3 levels may be increased in the CSF of heterozygous and homozygous *Grn^R493X^* mice at 12 months of age (Fig. 2D).

We found that heterozygous *Grn^R493X^* mice phenocopy heterozygous *Grn*^+/-^ mice reported in two previous studies [30,35]. Notably, they do not exhibit key features of human FTD-*GRN* including neuroinflammation and increased NfL levels in plasma or CSF. Nonetheless, like *Grn*^+/-^ mice, heterozygous *Grn^+/R493X^* mice have social and emotional deficits that recapitulate core symptoms of FTD-*GRN*. The presence of these behavioral deficits without observed increases in NfL levels suggests that in mice the social and emotional deficits may develop independently of overt neuronal death that is detectable by this sensitive biomarker. Taken together, although *Grn* gene dosage effects are also observed in mice, the heterozygous states manifest quite differently in humans and mice; the reasons for this species difference are unclear. One possibility that has been suggested is that the shorter lifespan of mice limits the disease progression [72]. It is also possible that genetic differences or species-specific differences in disease mediators may dampen the phenotype of heterozygous *Grn* mice. It is worth nothing that it is unlikely that the observed species difference are due to differences between the mouse and human progranulin proteins, as a recent study using a humanized *GRN* mouse model suggests that the mouse and human progranulin proteins are functionally equivalent in the context of preventing lipofuscinosis in the brain [73]. Overall, with the relatively mild phenotype of heterozygous *Grn*^+/-^ mice and *Grn^+/R493X^* mice, developing models of progranulin haploinsufficiency in larger animals may be warranted.

Overall, our findings in homozygous *Grn^R493X^* mouse brains are consistent with the phenotypes previously reported in homozygous *Grn* knockout mice. Key features of homozygous *Grn^R493X^* mouse from this study and from previous studies include neuroinflammation, microgliosis, astrogliosis, lipofuscin accumulation, vacuolation, upregulation of lysosomal genes, and TDP-43 pathology [27,28,37]. Notably, all of these phenotypes have also been observed in mouse models of NCL [30,74-77], raising the possibility that these observed phenotypes in homozygous *Grn* knockout and *Grn^R493X^* mice may reflect NCL rather than FTD-*GRN*. Our results support that, despite their more subtle phenotype, heterozygous *Grn*^+/-^ and *Grn^+/R493X^* mice are likely to be more appropriate models of FTD-*GRN* than the homozygous mice, as previously suggested [35,78].

Our study has limitations. Our analysis of neuroinflammation and lysosomal dysfunction relied heavilty on transcriptional changes; our studies assessing protein levels were more limited with smaller sample sizes. For our biomarker studies, it should be noted that the sample sizes were relatively small for measurements in CSF and in 18-month-old mice; therefore, these findings must be interpreted with caution. Additional studies are warranted, including measurements of C1qa and C3 levels in the CSF. Our behavioral studies were limited to male mice at 11 months of age; additional studies in female mice are necessary to address possible sex-specific phenotypes, which have been noted in the *Grn* knockout and *Grn^R493X^* mouse models [24,26,37,61].

In conclusion, we have performed in-depth characterization of the *Grn^R493X^* mouse model of FTD, including both heterozygous and homozygous mice. Overall, we found that heterozygous *Grn^R493X^* and homozygous knockin mice phenocopy their *Grn* knockout counterparts with respect to biochemical changes, neuroinflammation, levels of a plasma biomarker of neuronal damage, and behavioral and cognitive deficits. In contrast to human FTD-*GRN*, heterozygous *Grn^+/R493X^* and *Grn*^+/-^ mice do not show elevated levels of plasma NfL through 18 months of age, suggesting they do not have increased neuronal death compared to wild-type mice. These results may help to inform pre-clinical studies using this *Grn^R493X^* knockin model and other *Grn* knockout mouse models.

**Figure S1. *Grn^R493X^* mice have increased expression of several lysosomal genes in the thalamus.** Quantification of mRNA levels in the thalamus by qPCR (male and female mice). Bars represent mean ± SEM; the group sizes are indicated in each bar. * p < 0.05, ** p < 0.01, and **** p < 0.0001 compared to wild-type group, as determined by one-way ANOVA with Dunnett post hoc test. WT, wild-type; Het, *Grn^+/R493X^* heterozygous mice; KI, *Grn^R493X/R493X^* knockin mice.

**Figure S2. *Grn^R493X^* mice have changes in lysosomal proteins levels, inflammation, and TDP-43 levels in the cortex.** Additional western blot analysis of cortical tissues from 12-month-old mice (males and females, as indicated). WT, wild-type; Het, *Grn^+/R493X^* heterozygous mice; KI, *Grn^R493X/R493X^* knockin mice. These results are included in the quantification shown in Figures 1B, 2B, and 3A.

**Figure S3. Homozygous *Grn^R493X^* mice, but not heterozygous mice, have increased plasma NfL levels.** Plasma NfL levels at different ages (n = 3 per group) (A) and separated by sex (B) measured using the Quanterix Simoa platform. Bars represent mean ± SEM; the group sizes are indicated in each bar. * p < 0.05 and ** p < 0.01 compared to wild-type group, as determined by repeated measures two-way ANOVA with Tukey post hoc test (A) or one-way ANOVA with Dunnett post hoc test (B). WT, wild-type; Het, *Grn^+/R493X^* heterozygous mice; KI, *Grn^R493X/R493X^* knockin mice.

**Figure S4. Similar changes in plasma NfL levels detected with the Sigma SMC platform.** Plasma NfL levels at measured using the Sigma SMC platform. Bars represent mean ± SEM; the group sizes are indicated in each bar. * p < 0.05 and ** p < 0.01 compared to wild-type group, as determined by one-way ANOVA with Dunnett post hoc test. WT, wild-type; Het, *Grn^+/R493X^* heterozygous mice; KI, *Grn^R493X/R493X^* knockin mice; n.s., not significant.

**Figure S5. Homozygous *Grn* KO mice, but not heterozygous mice, have increased plasma NfL levels.** Plasma NfL levels at measured using the Quanterix Simoa platform. Bars represent mean ± SEM; the group sizes are indicated in each bar. * p < 0.05 compared to wild-type group, as determined by one-way ANOVA with Dunnett post hoc test. WT, wild-type; Het, *Grn^+/R493X^* heterozygous mice; KO, *Grn^–/–^* knockout mice.

**Figure S6. Plasma and CSF levels of GFAP in *Grn^R493X^* mice.** A) Plasma GFAP levels (male and female mice). B) CSF GFAP levels (female mice). Bars represent mean ± SEM; the group sizes are indicated in each bar. Statistical analysis was performed by one-way ANOVA with Dunnett post hoc test. WT, wild-type; Het, *Grn^+/R493X^* heterozygous mice; KI, *Grn^R493X/R493X^* knockin mice; n.s., not significant.

**Figure S7. *Grn^R493X^* mice show no differences in other behavioral tests performed.** Male mice, 11 months old. Bars represent mean ± SEM; the group sizes are indicated in each bar. Statistical analysis was performed by one-way ANOVA with Dunnett post hoc test. WT, wild-type; Het, *Grn^+/R493X^* heterozygous mice; KI, *Grn^R493X/R493X^* knockin mice.

## AUTHOR DECLARATIONS

### Ethics approval

All methods and animal procedures were approved by the Institutional Animal Care and Use Committee at Saint Louis University or the Harvard Medical Area Standing Committee on Animals.

### Consent to participate

Not applicable

### Consent for publication

Not applicable

### Availability of data and materials

All relevant data are within the paper and its Supplementary Information files.

### Authors’ contributions

D. M. Smith performed experiments, analyzed data, and prepared figures. G. Aggarwal performed experiments. M. L. Niehoff performed experiments. S. A. Jones performed experiments. S. Banerjee performed experiments. S. A. Farr performed experiments, analyzed data, and prepared figures. A. D. Nguyen designed the study, analyzed data, prepared figures, and wrote the manuscript. All authors read and approved the final manuscript.

## Acknowledgements

We thank Robert Farese, Jr., Tobias Walther, John Morley, and Thi Nguyen for helpful discussions, the Bluefield Project to Cure FTD for progranulin antibodies and *Grn* mouse plasma, Yuna Ayala for advice about TDP-43 analysis, Sami Barmada for qPCR primer sequences for sTDP-43 isoforms, the NeuroBehavior Laboratory at the Harvard NeuroDiscovery Center, the Quanterix Accelerator Lab, and Millipore Sigma for plasma NfL measurements using the SMC platform

