## Supplementary figures and images for "Biochemical, biomarker, and behavioral characterization of the *Grn^R493X^* mouse model of frontotemporal dementia"

### Fig. S1

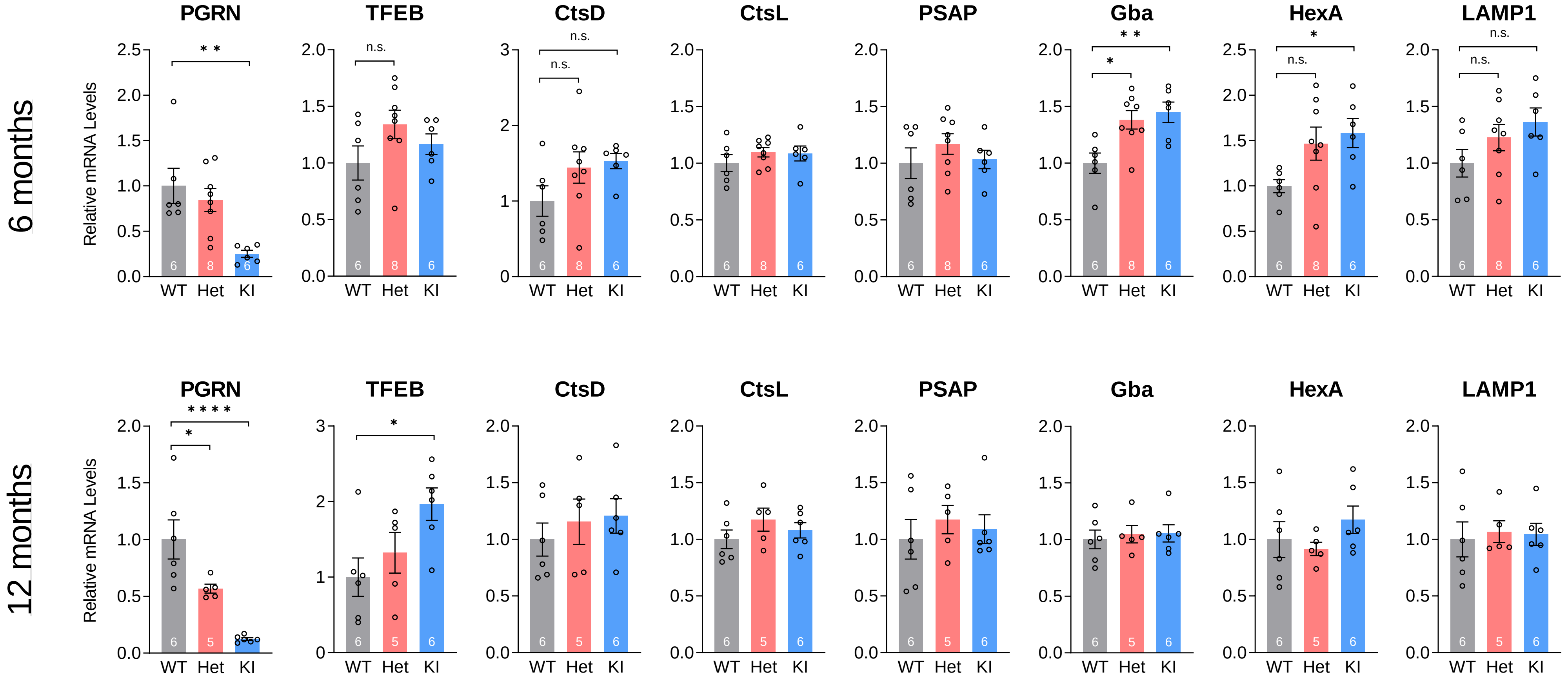

### Fig. S2

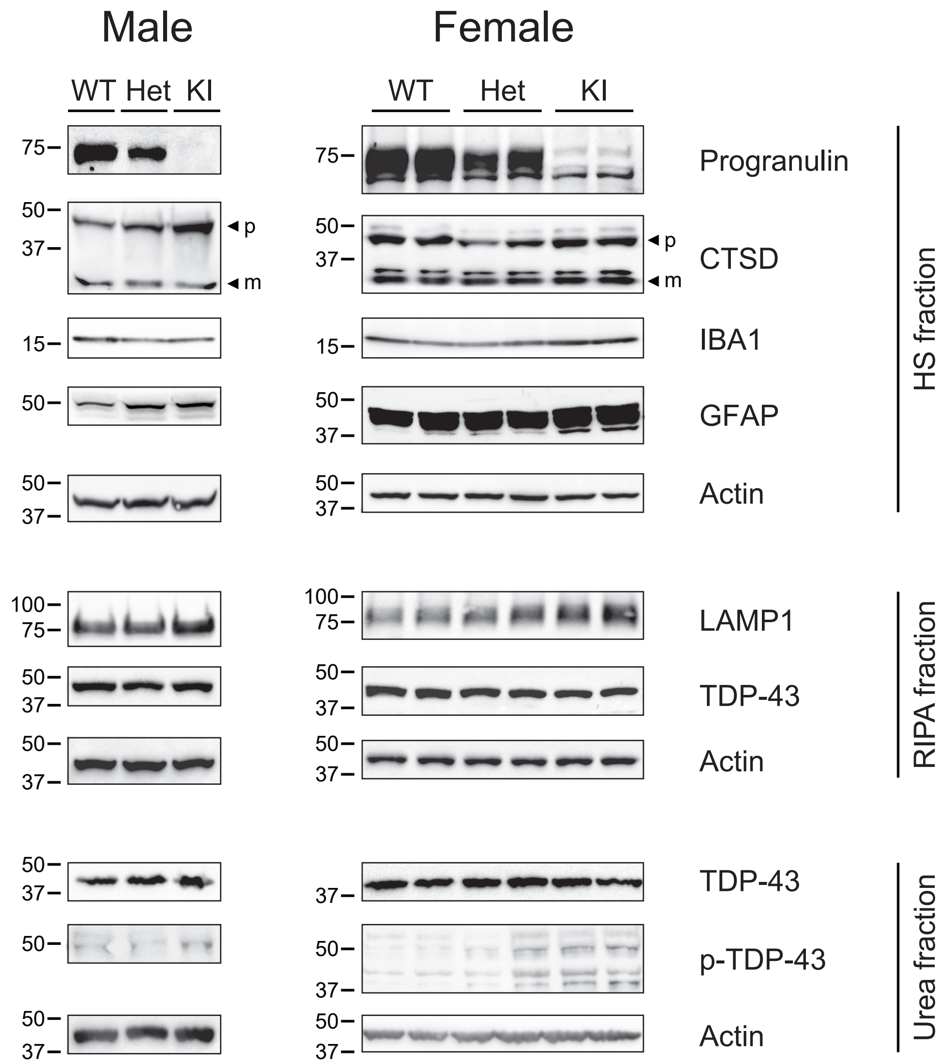

### Fig. S3

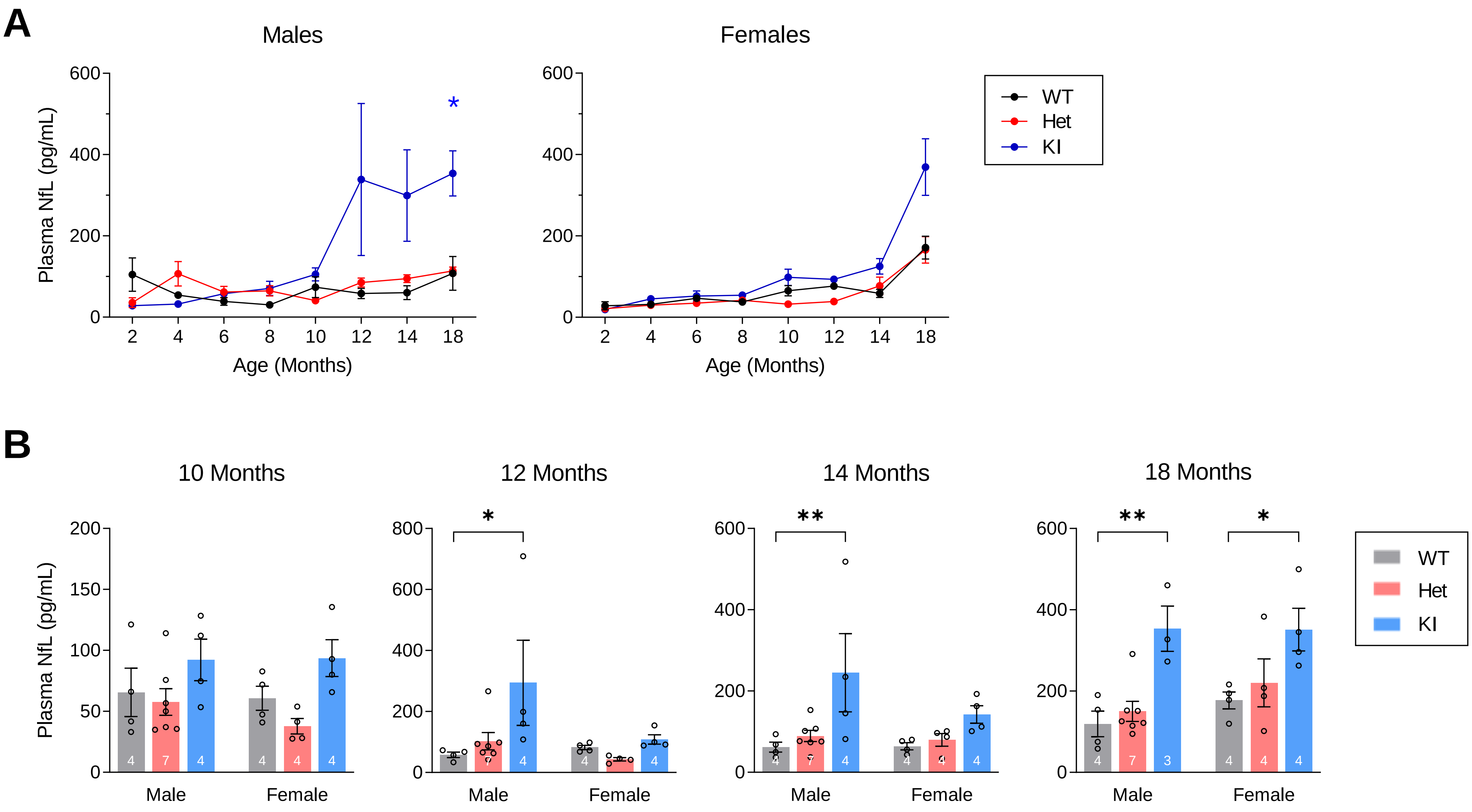

### Fig. S4

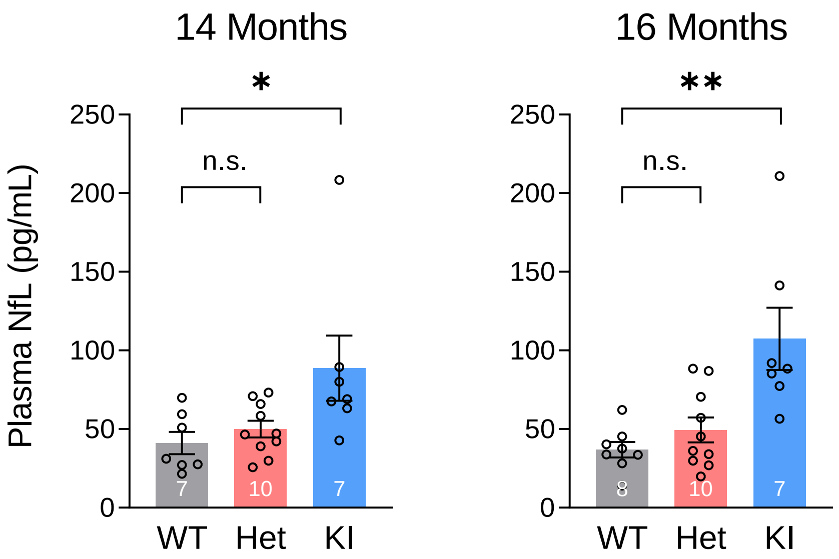

### Fig. S5

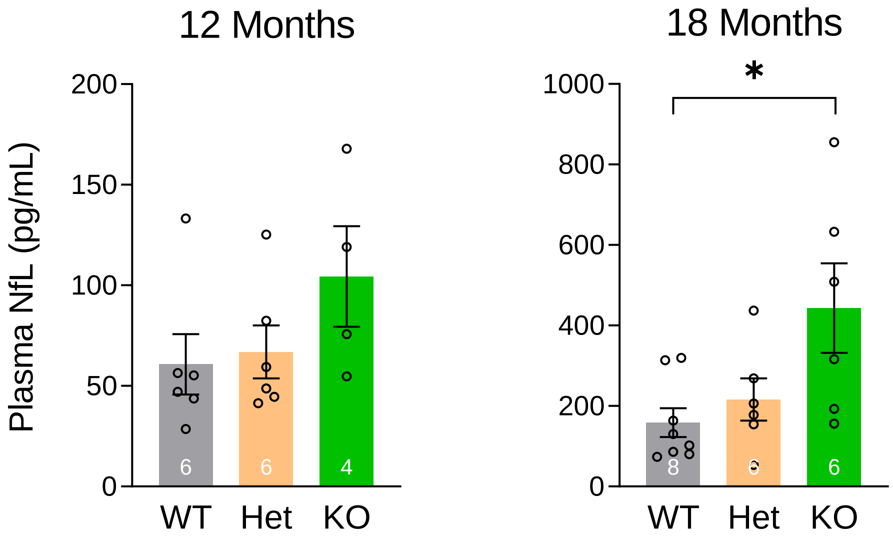

### Fig. S6

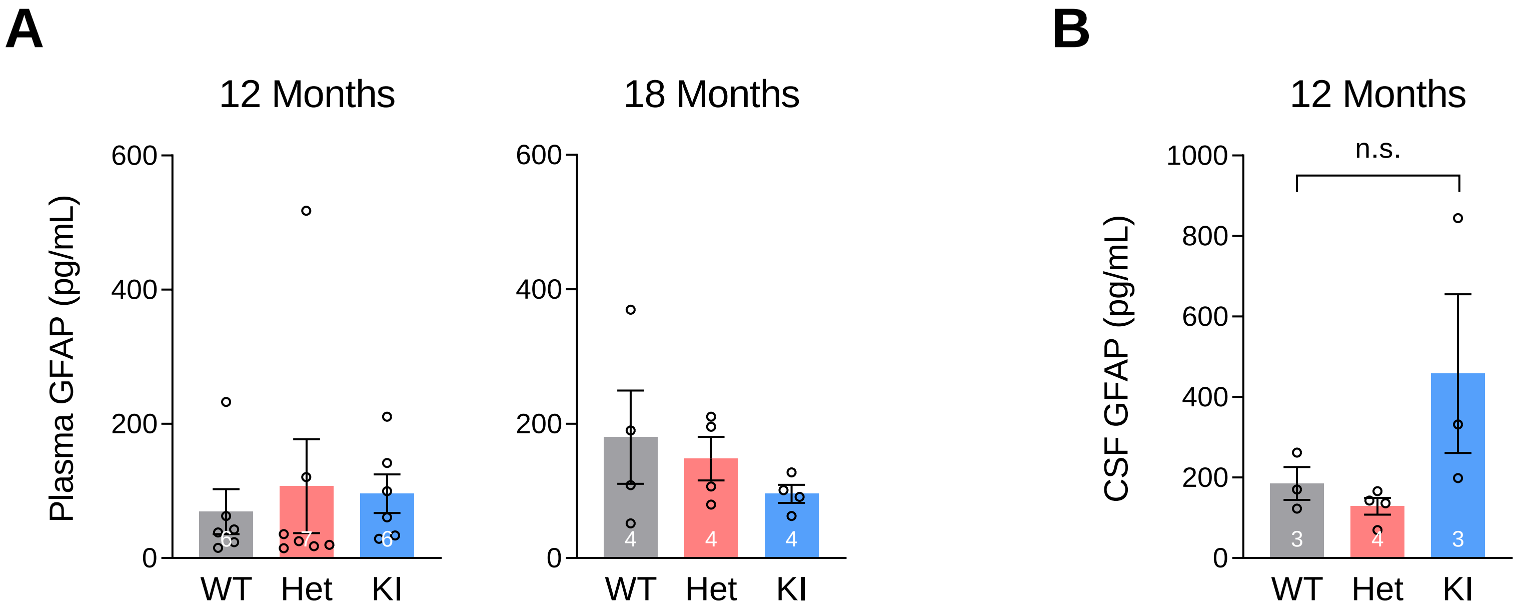

### Fig. S7

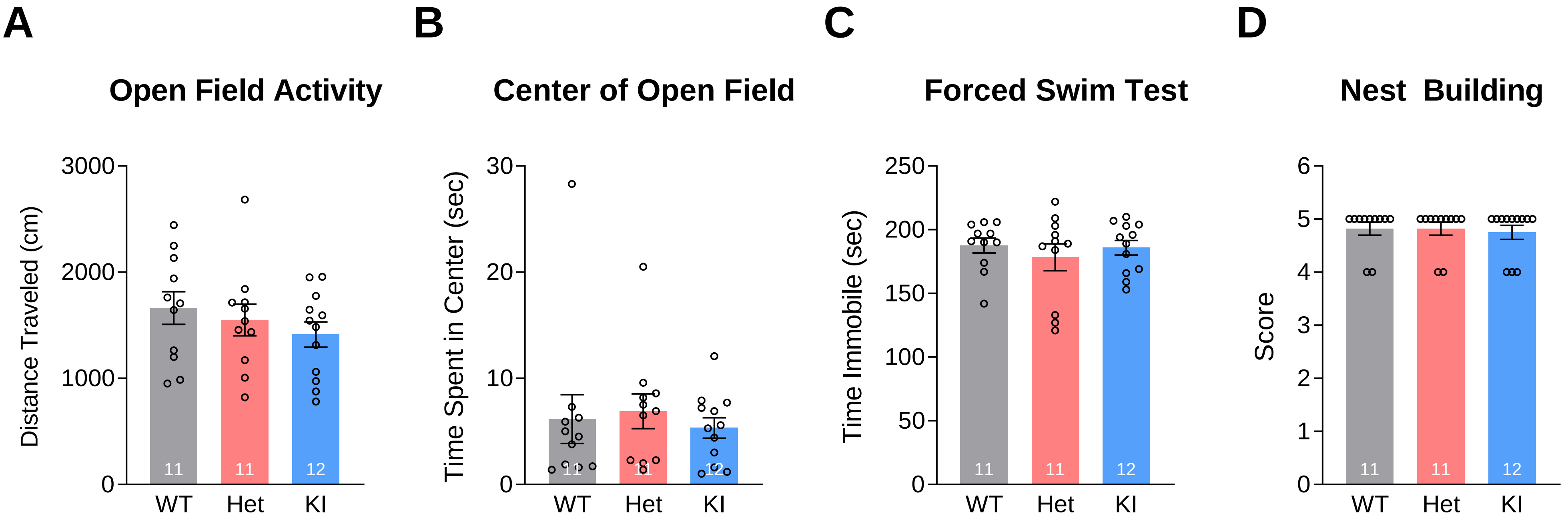
