## Supplementary material for "Biochemical, biomarker, and behavioral characterization of the *Grn^R493X^* mouse model of frontotemporal dementia": Table S1

**Table S1. qPCR primer sequences.**

| **Gene** | **Forward Primer** | **Reverse Primer** |
| --- | --- | --- |
| 36B4 | CACTGGTCTAGGACCCGAGAAG | GGTGCCTCTGAAGATTTTCG |
| Cyclo | GGCTCCGTCGTCTTCCTTTT | ACTCGTCCTACAGATTCATCTCC |
| PGRN | TGGTTCACACACGATGCGTTTCAC | AAAGGCAAAGACACTGCCCTGTTG |
| Tfeb | CCACCCCAGCCATCAACAC | CAGACAGATACTCCCGAACCTT |
| CtsD | CCTGGCTTCGTCCTCCTTC | GGCGATGACTGCATGGAGT |
| CtsL | ATCAAACCTTTAGTGCAGAGTGG | CTGTATTCCCCGTTGTGTAGC |
| PSAP | CCTGTCCAAGACCCGAAGAC | CAAGGAAGGGATTTCGCTGTG |
| Gba | GACCAACGCTTGCTGCTAC | ACAGCAATGCCATGAACGTA |
| Hexa | ACCTGGGAGGGGATGAAGT | ATGAAGGCCTGGATGTTGG |
| LAMP1 | GCCCACAAACCCCACTGTAT | TTTGGGCTGATGTTGAACGC |
| Iba1 | ATCAACAAGCAATTCCTCGATGA | CAGCATTCGCTTCAAGGACATA |
| GFAP | CGGAGACGCATCACCTCTG | AGGGAGTGGAGGAGTCATTCG |
| TNFα | ACGGCATGGATCTCAAAGAC | AGATAGCAAATCGGCTGACG |
| IL-1β | GCTTCAGGCAGGCAGTATC | AGGATGGGCTCTTCTTCAAAG |
| MCP1 | CTTCCTCCACCACCATGCA | CCAGCCGGCAACTGTGA |
| C1qa | AAAGGCAATCCAGGCAATATCA | TGGTTCTGGTATGGACTCTCC |
| C3 | CCAGCTCCCCATTAGCTCTG | GCACTTGCCTCTTTAGGAAGTC |
| TDP43 | AATCAGGGTGGGTTTGGTAACA | GCTGGGTTAATGCTAAAAGCAC |
| sTDP43-1 | AGAAGTGGAAGATTTGGTGTTCA | GCATGTAGACAGAAGTATTCCTATGG |
| sTDP43-2 | AGATTTGGTGGTAATCCAGTTCA | AGACAGAAGTATTCCTATGGCAGA |
| STMN2 | CAGAGGAGCGAAGAAAGTCTCA | CTAGATTAGCCTCACGGTTTTCC |
| UNC13A | CAGGGTCGGCTTGATTCTG | GACTCTTGCTTGTCACCTGTG |
